# Isolation and Characterization of a Roseophage Representing a Novel Genus in the N4-like Rhodovirinae Subfamily Distributed in Estuarine Waters

**DOI:** 10.1101/2024.10.08.617335

**Authors:** Xingyu Huang, Chen Yu, Longfei Lu

**Affiliations:** Institute of Tibetan Plateau Research, Chinese Academy of Sciences,. No. 16 Lincui Road, Chaoyang District, Beijing 100101, P.R. China; State Key Laboratory of Marine Environmental Science, College of Ocean & Earth Sciences, Fujian Key Laboratory of Marine Carbon Sequestration, Xiamen University, Xiamen 361102, PR China; Key Laboratory of Tropical Marine Ecosystem and Bioresource, Fourth Institute of Oceanography, Ministry of Natural Resources, Beihai 536000, China

**Keywords:** Bacteriophage, Roseophage, Isolation, Phylogenetic analysis, Auxiliary metabolism gene, Biogeography

## Abstract

**Background:** *Roseobacteraceae*, often referred to as the marine roseobacter clade (MRC), are pivotal constituents of bacterial communities in coastal and pelagic marine environments. During the past two decades, 75 roseophages that infect various *Roseobacteraceae* lineages have been isolated. The N4-like *Rhodovirinae* subfamily, which encompasses 15 members, represents the largest clade among these roseophages.

**Results:** In this study, we isolated a novel roseophage, vB_DshP-R7L, that infects *Dinoroseobacter shibae* DFL12 from Xiamen Bay in the East China Sea. Conserved genes of *Schitoviridae* have been identified in the genome of vB_DshP-R7L, and following phylogenetic analysis suggest that the newly isolated phage is a member of the *Rhodovirinae* subfamily and is indicative of a newly proposed genus, *Xianganvirus*. The genome of vB_DshP-R7L harbors six auxiliary metabolic genes (AMGs), most of which potentially enhance DNA *de novo* synthesis. Additionally, a gene encoding ribosomal protein was identified. Comparative genomic analysis of AMG content among *Rhodovirinae* indicates a distinct evolutionary history characterized by independent ancient horizontal gene transfer events. Read-mapping analysis reveals the prevalence of vB_DshP-R7L and other *Rhodovirinae* roseophages in estuarine waters.

**Conclusions:** Our work illustrates the genomic features of a novel roseophage clade among N4-like *Rhodovirinae*. The AMG content of vB_DshP-R7L are under severe purification selection, which revealed their possible ecological importance. We also demonstrated that vB_DshP-R7L and other *Rhodovirinae* roseophages are restrictively distributed in estuaries. Our isolation and characterization of this novel phage expand the understanding of the phylogeny, gene transfer history and biogeography of N4-like *Rhodovirinae* infecting marine *Roseobacteraceae*.

## Background

Phages are viruses that can infect bacteria, and play critical roles in biogeochemical and ecological functions [1]. In the oceans, phages outnumbering their hosts by an order of magnitude [1, 2], influencing microbial community dynamics by modulating host mortality and facilitating horizontal gene transfer [1, 3]. Marine phages frequently contain auxiliary metabolic genes (AMGs). AMGs are viral genes originally acquired from host genomes through horizontal gene transfer, enable viruses to manipulate host cellular machinery during infection, alleviate metabolic bottlenecks, and augment host metabolism to maximize viral production [4–7].

The marine roseobacter clade (MRC) represents one of the most abundant heterotrophic bacterial taxa in coastal ecosystems, comprising approximately 5% and 20% of bacterial communities in open oceans and nearshore ecosystems [8, 9]. All members of the MRC form a single monophyletic group, classified as family *Roseobacteraceae*, which is involved in multiple metabolic pathways [9, 10]. The *Roseobacteraceae* are pivotal in marine biogeochemical cycles due to their roles in both aerobic and anaerobic photosynthesis, and they are the primary reducers of dimethylsulfoniopropionate (DMSP) produced by dinoflagellates [10–12].

To date, 75 roseophages that infect members of the *Roseobacteraceae* have been identified [13–37] (Table 1). Of these, two belong to *Monodnaviria*, with the others classified as *Caudoviricetes*. Comparative genomic analyses have delineated five major clades among the *Caudoviricetes* roseophages, namely *Schitoviridae*, *Cobavirus*, *Autographiviridae*, Cbk-like, and Chi-like clades [14, 18, 38, 39]. Roseophages from the *Cobavirus*, *Autographiviridae*, and Cbk-like clades have shown a global distribution [13, 14, 18].

**Table 1.**
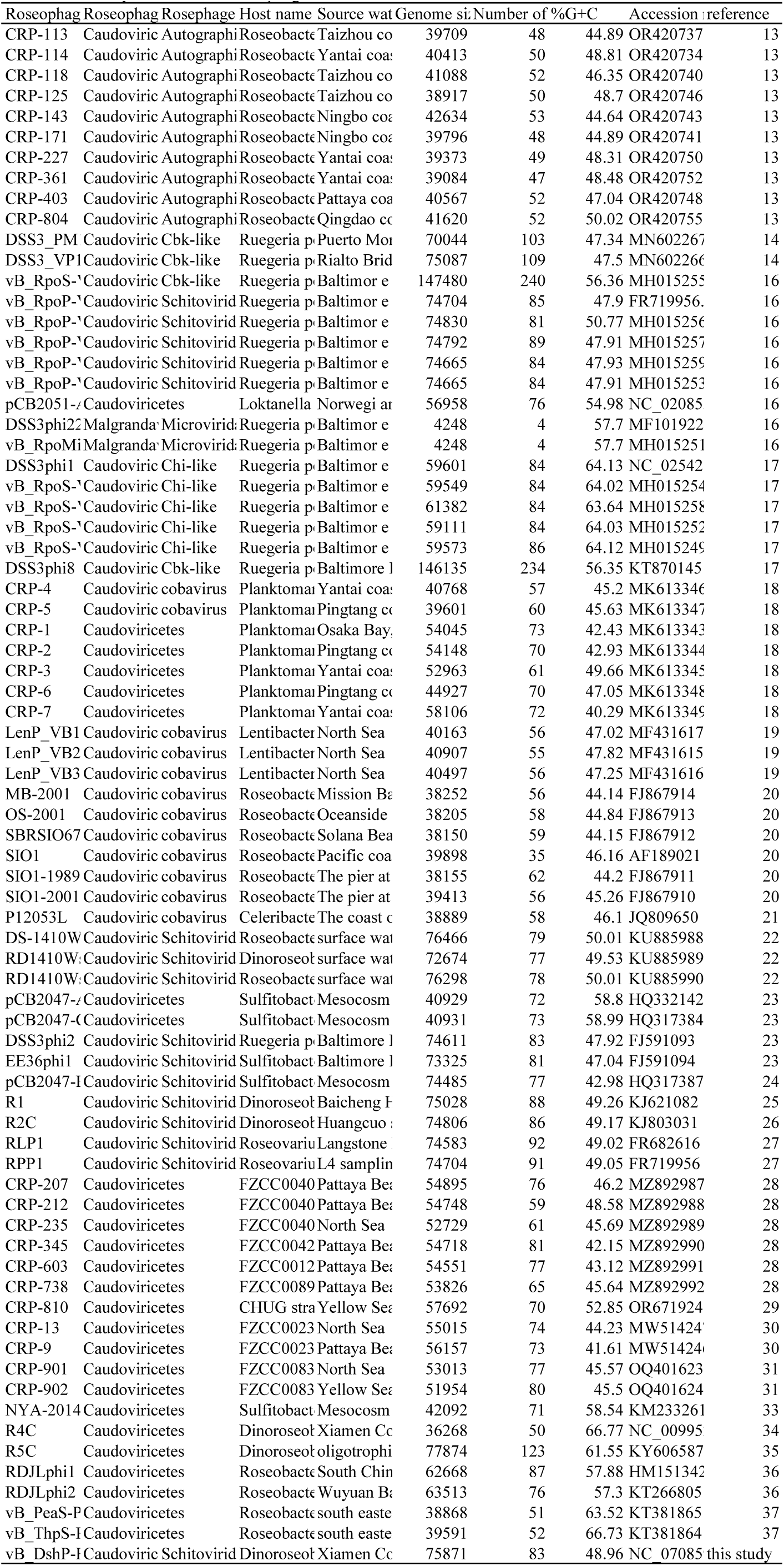
Summary of all isolated roseophages.

Roseophages from the *Schitoviridae* clade constitute the largest group of isolated roseophages. The family *Schitoviridae*, previously recognized as the N4-like phage clade, share several conserved genes forming a universal backbone, notably three DNA-dependent RNA polymerase genes [40, 41]. The genomes of *Schitoviridae* are transcribed in three programmed stages during infection, with each RNA polymerase functioning at a specific stage [41]. All roseophages within the *Schitoviridae* form a unique monophyletic group classified as the subfamily *Rhodovirinae*, which is further divided into seven genera: *Aorunvirus*, *Raunefjordvirus*, *Aoquinvirus*, *Pomeroyivirus*, *Sanyabayvirus*, *Plymouthvirus*, and *Baltimorevirus* [40]. Comparative genomic analysis indicates that diverse AMGs present in *Rhodovirinae* genomes, which highlights their potential significant ecological roles [39]. These AMGs predominantly involved in the DNA *de novo* synthesis pathway [39]. Despite detailed genomic characterizations, the distribution of *Rhodovirinae* remains poorly understood [27].

In this study, we report a novel roseophage, vB_DshP-R7L, infecting *Dinoroseobacter shibae* DFL12 isolated from the coastal waters of Xiamen. Phylogenetic analysis suggests that vB_DshP-R7L represents a previously unrecognized genus within the N4-like *Rhodovirinae* subfamily. Six AMGs were identified in its genome, and their evolutionary history was comprehensively analyzed. The tRNA content in the vB_DshP-R7L genome may enhance the expression of its AMGs. Furthermore, metagenomic analysis revealed the prevalence of vB_DshP-R7L and its *Rhodovirinae* relatives in estuarine waters.

## Methods

### Isolation and purification of vB_DshP-R7L

vB_DshP-R7L was isolated from the surface water of Xiamen Bay station S03 (118.03N, 24.43E) during the Xiamen nearshore cruise in November 2013. Seawater samples were filtered through a 0.22 µm membrane and stored at 4°C in darkness. For cultivation, 5 mL of *Dinoroseobacter shibae* DFL12 host culture was incubated at 20°C and agitated at 100 rpm in RO medium (1 g·L^−1^ yeast extract, 1 g·L^−1^ peptone, 1 g·L^−1^ sodium acetate, 1 L artificial seawater, with a pH maintained between 7.4 and 7.8) until reaching the exponential growth phase. 1 ml of the preserved seawater filtrate was then introduced to this culture medium and incubated for 7 days. The phage was isolated and purified through five successive rounds using the double-layer agar plate method.

After purification, phage plaques were collected with a pipette tip and expanded in 1 L of liquid RO medium containing *Dinoroseobacter shibae* DFL12 in its exponential phase. The culture was subsequently centrifuged at 12000×g for 10 minutes at 4 °C. The supernatant was transferred to a fresh container and the phage particles were concentrated via overnight precipitation using 100 g of PEG8000, followed by centrifugation at 10000×g for 60 minutes at 4 °C. The resulting precipitate was resuspended in 6 ml of SM buffer (100 mM NaCl, 8 mM MgSO_4_, 50 mM Tris-HCl) and the phage-containing band was further purified using 30 kDa super-filters (UFC503096, Millipore) and adjusted to the desired concentration with SM buffer following previously reported method [35].

The purified viruses were also tested against various roseophage hosts including *Erythrobacter litoralis* DSM 8509, *Erythrobacter* sp. JL475, *Erythrobacter* DSM 6997, and *Roseobacter denitrificans* DSM 7001, all cultured to their exponential phases. The host range was determined using the double-layer agar plate method with each bacterial strain tested in triplicate.

### Transmission electron microscopy (TEM) and one-step growth curve

For electron microscopy, the purified and desalted phage particles were adsorbed onto 200-mesh carbon-coated copper grids for 10–30 minutes in the dark. After staining with 1% phosphotungstic acid for 30 minutes, the samples were examined using a JEM-2100 transmission electron microscope (JEOL, Tokyo, Japan) at an 80 kV voltage. Images were captured using an HPF-TEM system (Gatan Inc., Pleasanton, CA, USA).

A one-step growth curve assay was conducted to evaluate the infectivity and replication capability of vB_DshP-R7L. Phages were added to 200 ml culture of *Dinoroseobacter shibae* DFL12 in its exponential phase at a concentration of 1×10^6^ ml^−1^ and incubated at 28 °C with continuous shaking. Samples were collected every 30 minutes, and viral abundance was quantified using the epifluorescence microscopy method [42].

### Characterization of structural protein

Structural proteins of vB_DshP-R7L were analyzed by mixing phage particles with loading buffer and heating to 100°C for 5 minutes. The heated samples were separated using 12% sodium dodecyl sulfate-polyacrylamide gel electrophoresis (SDS-PAGE) and the gels were stained with silver. Protein analysis was conducted using liquid chromatography–mass spectrometry at the Shanghai Applied Protein Technology.

### DNA extraction and genome annotation

Phage DNA was extracted using the phenol-chloroform method, following the protocols described by Yang [35]. The extracted DNA was sequenced using an Illumina HiSeq 2500 sequencer with a paired-end, 2 × 251 bp configuration. The sequencing produced a total of 1.56 gigabases of clean reads. The CLC Genomics Workbench software was utilized to analyze the high-throughput sequencing data and to assemble these reads into a complete genome sequence. This sequencing and assembly process was performed by MAGIGENE Corporation. The complete genome sequence has been deposited in GenBank under the accession number MZ773648.1.

The genomic annotation was carried out using Prodigal v2.6.3 [43] to predict putative open reading frames (ORFs) within the vB_DshP-R7L genome. Functional annotation of these viral ORFs was performed using the DIAMOND BLASTp tool against the NCBI non-redundant protein sequence database, updated as of January 8, 2024. Additionally, InterProScan v5.52-86.0 was employed to annotate viral ORFs, referencing the Pfam-A database [44]. tRNAscan-SE was utilized to identify tRNAs within the genome, accepting only results classified as high-quality [45]. The involvement of auxiliary metabolic genes in cellular metabolic pathways was determined manually by analyzing ORF prediction and BLAST results against the KEGG pathway database [46].

### Comparative genomics and phylogenetic analysis

The genome structure of vB_DshP-R7L and its homologous phages was compared using the EasyFig tool, which employs the TBLASTX comparison method [47]. Genomes of all isolated roseophages and *Schitoviridae* phages were retrieved from GenBank on September 1, 2024, and from a recent study [48]. Viral proteins were clustered at a 70% identity threshold using CD-hit, adhering to a previously established method [49]. Sixteen conserved genes shared among the *Schitoviridae* phages were identified in each viral genome. These conserved genes were concatenated and used to construct the phylogenetic tree, along with marker genes encoding the virion-encapsidated RNA polymerase and the major capsid protein to construct separate phylogenetic trees.

The phylogenetic tree of all isolated roseophages genomes was generated using VIPtree software [50]. Concatenated conserved genes and marker genes were aligned using MAFFT (v.7.453) under default settings and subsequently trimmed with trimAL (v.1.4) to eliminate blank regions. The aligned sequences were used to construct a maximum likelihood phylogenetic tree with Raxml-ng [18], employing the LG+G8+F model and starting with an initial tree configuration of 10. The resultant tree was visualized using iTOL [51]. Phylogenetic trees of AMGs in the vB_DshP-vB_DshP-R7L and their homologs were also constructed and visualized using the same method. Additionally, amino acid sequence comparisons of the ribosomal protein encoding AMG and its homologs were visualized by AliView [52].

The ratio of synonymous to nonsynonymous substitutions (dN/dS) was employed to examine the genetic variation and evolutionary patterns of vB_DshP-R7L and its homologs. The substitution rates were calculated using the branch-site unrestricted statistical test for episodic diversification available in the HyPhy package [53]. In this analysis, genes from vB_DshP-R7L served as the test group, and their homologs formed the background group.

The tRNA Adaptation Index (tAI) was used to measure the tRNA usage preferences of each gene. This index calculates the tRNA copy number and the wobble interaction between the codon and anticodon using stAIcalc [54]. The tAI value of a gene, ranging from 0 to 1, is determined by the geometric mean of the normalized codon usage values. A higher tAI value indicates greater availability and translatability from the tRNA pool, signifying a gene’s efficient translation.

### Distribution of the *Rhodovirinae* roseophages in the global ocean

A comprehensive assessment of the relative abundance of isolated *Rhodovirinae* roseophages was conducted using a total of 294 marine metagenomic datasets. These datasets were from Global Ocean Viromes 2.0 [55], South China Sea [56, 57], East China Sea [58], Pearl River estuary virome [59], Chesapeake bay [60], Delaware Bay [60], Goseong bay [61] and Yangshan harbour [62] (Table S1). Marine metagenomic reads were mapped against the isolated *Rhodovirinae* roseophages genomes using CoverM v0.7.0 with previously with a previously reported method (--min-read-percent-identity 95 --min-read-aligned-length 50) [29]. The relative abundances of the phages were normalized to mapped read counts per kilobase pair of genome per million read counts (RPKM). Phages were considered present at a site if their genome coverage exceeded 30% in the dataset. The distribution of *Rhodovirinae* roseophages across global oceans was visualized using the R ggmap package.

## Results

### Virion shape, host range and one-step growth curve of vB_DshP-R7L

vB_DshP-R7L was isolated from the surface waters of Xiamen Bay at the Xiangan coastal station S03 (118.03N, 24.43E) in November 2013. Electron microscopy revealed that the vB_DshP-R7L virions possess an isometric head approximately 70 nm in diameter and a short tail about 40 nm long (Figure. 1a). Host range assessments conducted on other hosts of *Rhodovirinae* roseophages including *Erythrobacter litoralis* DSM 8509, *Erythrobacter* sp. JL475, *Erythrobacter* DSM 6997, *Roseovarius nubinhibens*, *Roseovarius* sp. 217, *Ruegeria pomeroyi* and *Roseobacter denitrificans* DSM 7001 demonstrated that vB_DshP-R7L has a limited host range, failing to infect reported hosts of *Rhodovirinae*. Further genomic comparisons of vB_DshP-R7L against CRISPR spacers within the *Roseobacteraceae* and the IMG/VR environmental CRISPR spacer database indicated exclusive matches with spacers from *Dinoroseobacter shibae* DFL12.

**Fig. 1.**
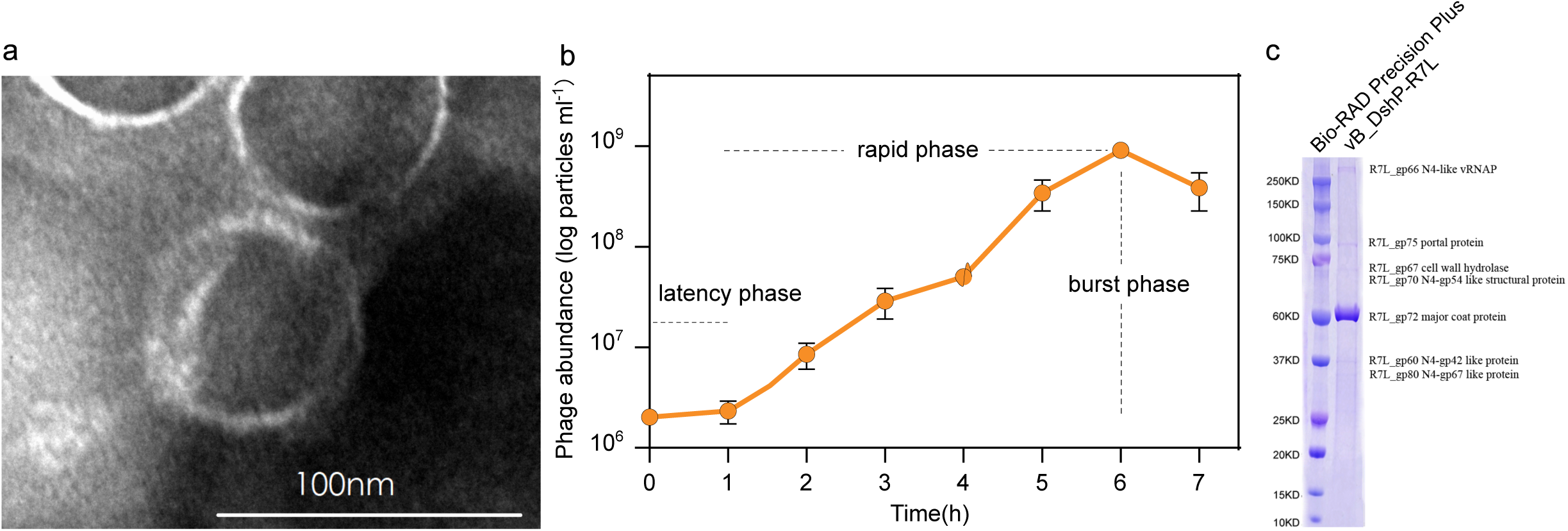
Characterization of vB_DshP-R7L Phage. a)Transmission electron micrograph of vB_DshP-R7L particles. b)One-step growth curves of vB_DshP-R7L. Latent period and burst size of vB_DshP-R7L phage were 1h and 109 particles per milliliter, respectively. C)Identification of structural proteins of vB_DshP-R7L phage. Purified virus particles were subjected to 8%-18% protein SDS-polyacrylamide gel electrophoresis and bands were labeled by protein molecular weight and mass spectrometry results.

The one-step growth curve of vB_DshP-R7L revealed a latency period of approximately 1 hour, followed by a rapid lysis phase lasting 5 hours and achieving a peak viral density of 1e^-9^/ml (Figure. 1b). The observed burst size was 320 particles per infected cell, significantly smaller than typical *Rhodovirinae* burst sizes, which often exceed 1500 [23] and far less than the 3,000 particles produced by *Enquatrovirus* N4 [63].

### Genome content and phage structural proteins

vB_DshP-R7L is a linear dsDNA phage with a genome length of 75.87 kb and a G+C content of 48.96%. A 99 bp repeat sequence was identified at the 3’ end of the genome. Analysis of the genome revealed 84 open reading frames (ORFs), most of which encode hypothetical proteins. 25 homologous genes shared with other *Rhodovirinae* roseophages were identified in the genome of vB_DshP-R7L (Table 2). The genome also encodes two tRNAs for proline and isoleucine.

**Table 2.**
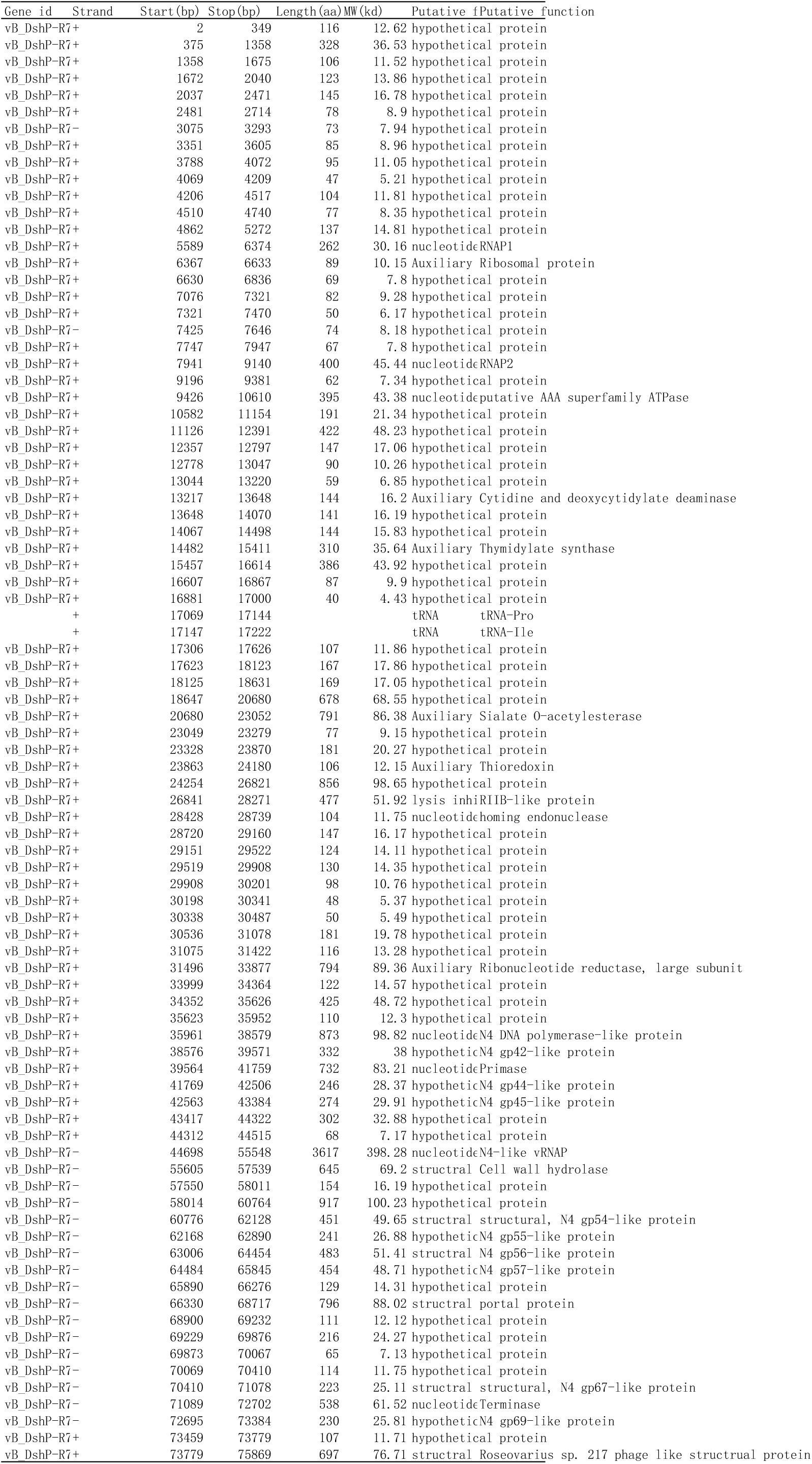
Genome annotation of vB_DshP-R7L.

Structural protein analysis through gradient SDS-PAGE electrophoresis and mass spectrometry identified five conserved structural proteins in *Schitoviridae* (Figure. 1c). R13LH_gp66 (398.28 kDa), was predicted to function as an N4-like viral RNA polymerase, and has the largest molecular weight among all structural proteins. The most abundant structural protein, based on peptide counts from mass spectrometry, was vB_DshP-R7L_gp72, predicted as the N4-like major capsid protein. vB_DshP-R7L_gp67, the second most abundant, was predicted as a cell wall hydrolase. vB_DshP-R7L_gp70 and vB_DshP-R7L_gp75 were identified as homologs of the N4 gp54-like structural protein and the N4-like viral portal protein, respectively.

### Phylogenetic analysis of vB_DshP-R7L

To clarify the taxonomy of vB_DshP-R7L, we conducted multiple phylogenetic analyses based on amino acid sequences. A proteomic tree of all isolated roseophages constructed with ViPTree indicates that vB_DshP-R7L forms a separate branch within the *Schitoviridae* family (Table 1) (Figure 2a). For classification within the *Schitoviridae*, we utilized 16 concatenated conserved genes previously reported, which are homologous to *Enterobacteria* phage N4 genes gp16, gp24, gp39, gp42-45, gp52, gp54-59 and gp69 [48]. These genes, found in over 90% of *Schitoviridae* phages, are conserved across both isolated and metagenomically derived genomes [48]. The concatenated conserved gene tree of vB_DshP-R7L and all subfamilies of *Schitoviridae* positions this newly isolated roseophage as a novel clade within the subfamily *Rhodovirinae* (Figure 2b). The distance between vB_DshP-R7L and its closest relative, *Baltimorevirus*, is 0.1053, which is similar to the distance between *Aorunvirus* and *Plymouthvirus* within *Rhodovirinae* (0.1048). This suggests that vB_DshP-R7L represent a novel genus within subfamily *Rhodovirinae,* family *Schitoviridae*.

**Fig. 2.**
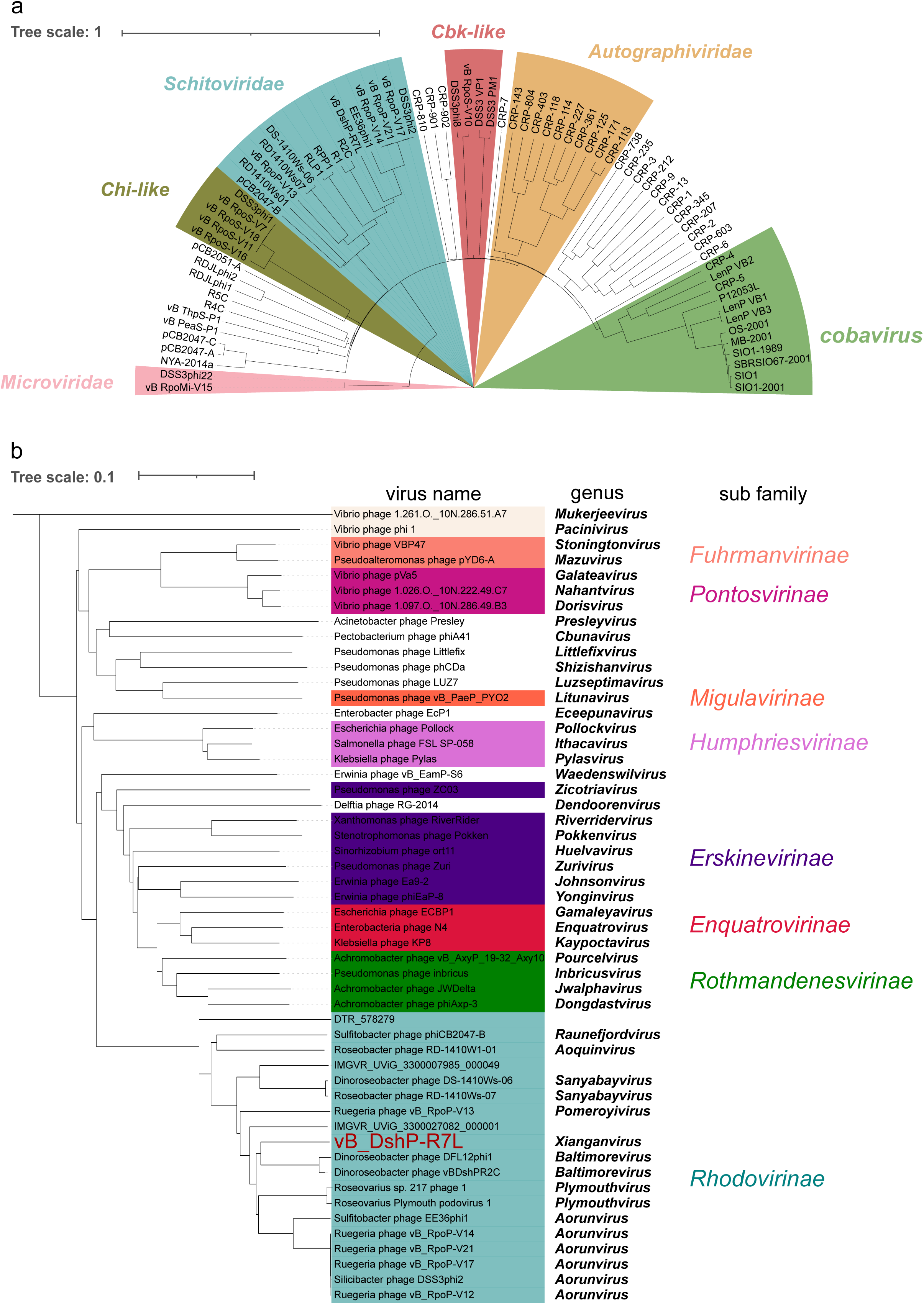
Phylogenetic relationships of vB_DshP-R7L and related phages a)Phylogenetic analyses of protein content similarity of all isolated roseophages using VipTree. The background color of six reported roseophage clades on the tree consistently with their labels. b)Phylogenetic tree based on 16 concatenate conserved genes of *Rhodovirinae* and their relatives in *Schitoviridae*. The phage vB_DshP-R7L, isolated in this study, is highlighted in red. The background color of each subfamily on the *Schitoviridae* tree corresponds consistently with their labels.

Additional evidence supporting this classification was obtained by constructing phylogenetic trees based on two marker genes of *Schitoviridae*, which are homologous to *Enterobacteria* phage N4 gp16 and gp56, which encodes vRNAP2 (Virion-encapsidated RNA polymerase) and MCP (Major capsid protein) (Figure S1). The distances between vB_DshP-R7L and *Baltimorevirus* (0.14 and 0.05) exceed the distance between *Aorunvirus* and *Plymouth* within *Rhodovirinae* (0.08 and 0.01). Clustering of all *Schitoviridae* phages by their shared genes using vContact2.0 marked vB_DshP-R7L as a unique genus-level viral cluster (VC), further substantiating its status as a distinct genus.

Comparative genomic analysis of vB_DshP-R7L with its closely related genera, *Baltimorevirus* and *Plymouthvirus*, along with the representative phage of *Schitoviridae*, *Enterobacteria* phage N4, revealed a conserved gene organization (Figure 3).

**Fig. 3.**
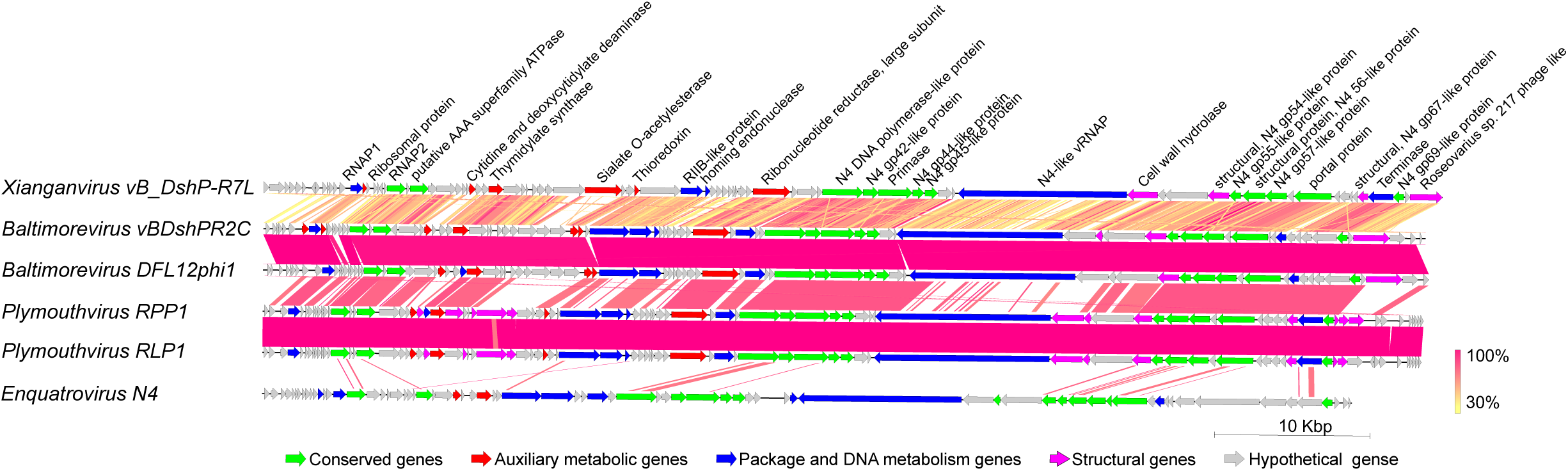
Genome comparison of the vB_DshP-R7L with its closest relatives and *Enquatrovirus* N4 using EasyFig. The open reading frames are indicated by leftward or rightward arrows depending on the direction of transcription. The colors of the arrows represent the predicted gene function, and homologs are linked according to the similarity values obtained by tblastx comparison. The color gradient representing similarity levels progresses from yellow to red, as detailed in the legend.

### Biogeography of vB_DshP-R7L and N4-like *Rhodovirinae* roseophages

To investigate the ecological distribution of vB_DshP-R7L and other *Rhodovirinae* roseophages, we conducted a metagenomic read-mapping analysis using 294 marine metagenomic datasets from both open oceans and estuaries as detailed in Methods. Our analysis revealed that all *Rhodovirinae* roseophages were isolated exclusively in estuarine environments. Specifically, phages R1, R2C, and vB_DshP-R7L were detected in metagenomic samples from the Pearl River Estuary, whereas vB_RpoP-V12 was found in metagenomic samples from Delaware Bay (Figure 4).

**Fig. 4.**
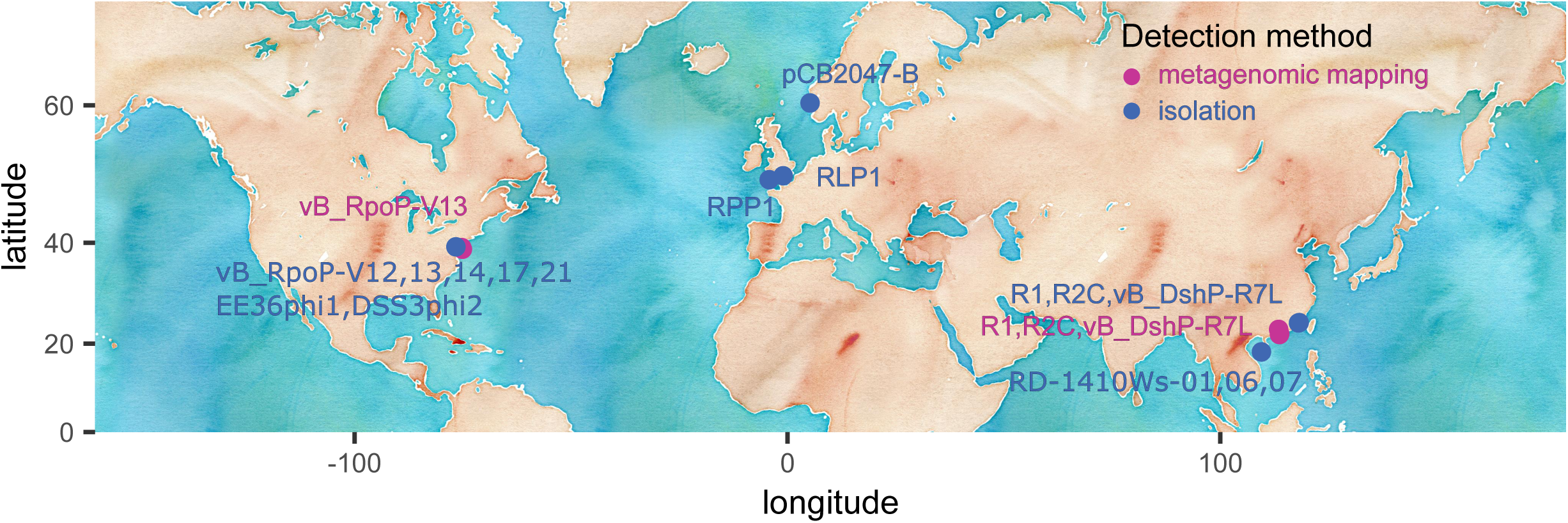
Global distribution of *Rhodovirinae* roseophages in marine waters. Stations where *Rhodovirinae* roseophages have been isolated are marked with blue circles, and those where they have been detected in metagenomes are marked with purple circles. The names of the *Rhodovirinae* roseophages are labeled on the map near the stations where they were detected. Each name is colored according to the method of detection.

### Function of phage auxiliary metabolic genes

We identified six auxiliary metabolic genes (AMGs) in the genome of vB_DshP-R7L, these include vB_DshP-R7L_gp29 (*dcd*) encoding dCMP deaminase, vB_DshP-R7L_gp32 (*thyX*) encoding thymidylate synthase, vB_DshP-R7L_gp43 (*trx*) encoding thioredoxin, vB_DshP-R7L_gp55 (*rnr*) encoding ribonucleotide reductase, vB_DshP-R7L_gp15 encoding a ribosomal protein, and vB_DshP-R7L_gp40 (*nanS*) encoding sialate O-acetylesterase.

In *Rhodovirinae*, the *trx* and *rnr* genes are conserved across all genomes, while the *dcd* and *thyX* genes are also prevalent [39]. Thioredoxin reductase, encoded by the *trx* gene, functions as a hydrogen donor, utilizing NADPH to transfer electrons to the ribonucleotide reductase encoded by the *rnr* gene, thereby reducing ribonucleoside diphosphate (rNDP) to deoxyribonucleoside diphosphate (dNDP) [64, 65]. DCMP deaminase, encoded by the *dcd* gene, deaminates deoxycytidylic acid (dCMP) to deoxyuridine monophosphate (dUMP). Thymidylate synthase, encoded by the *thyX* gene, is an essential enzyme for pyrimidine synthesis, converting dUMP to deoxythymidine monophosphate (dTMP) [66]. These four AMGs are related to the DNA *de novo* synthesis pathway and may enhance the production of precursors required for DNA synthesis, thus supporting DNA replication and modifications during infection (Figure 5). vB_DshP-R7L_gp15 encode a ribosomal protein bl12, which forms the prominent stalk-like structure on the large ribosomal subunit in bacteria and is the most common ribosomal protein encoded by viruses from aquatic environments [67]. vB_DshP-R7L_gp40 is predicted as *nanS* gene which encodes sialate O-acetylesterase. This enzyme catalyzes the hydrolysis of N-acetylneuraminic acid, a compound commonly found in the genomes of pathogenic bacteria.

**Fig. 5.**
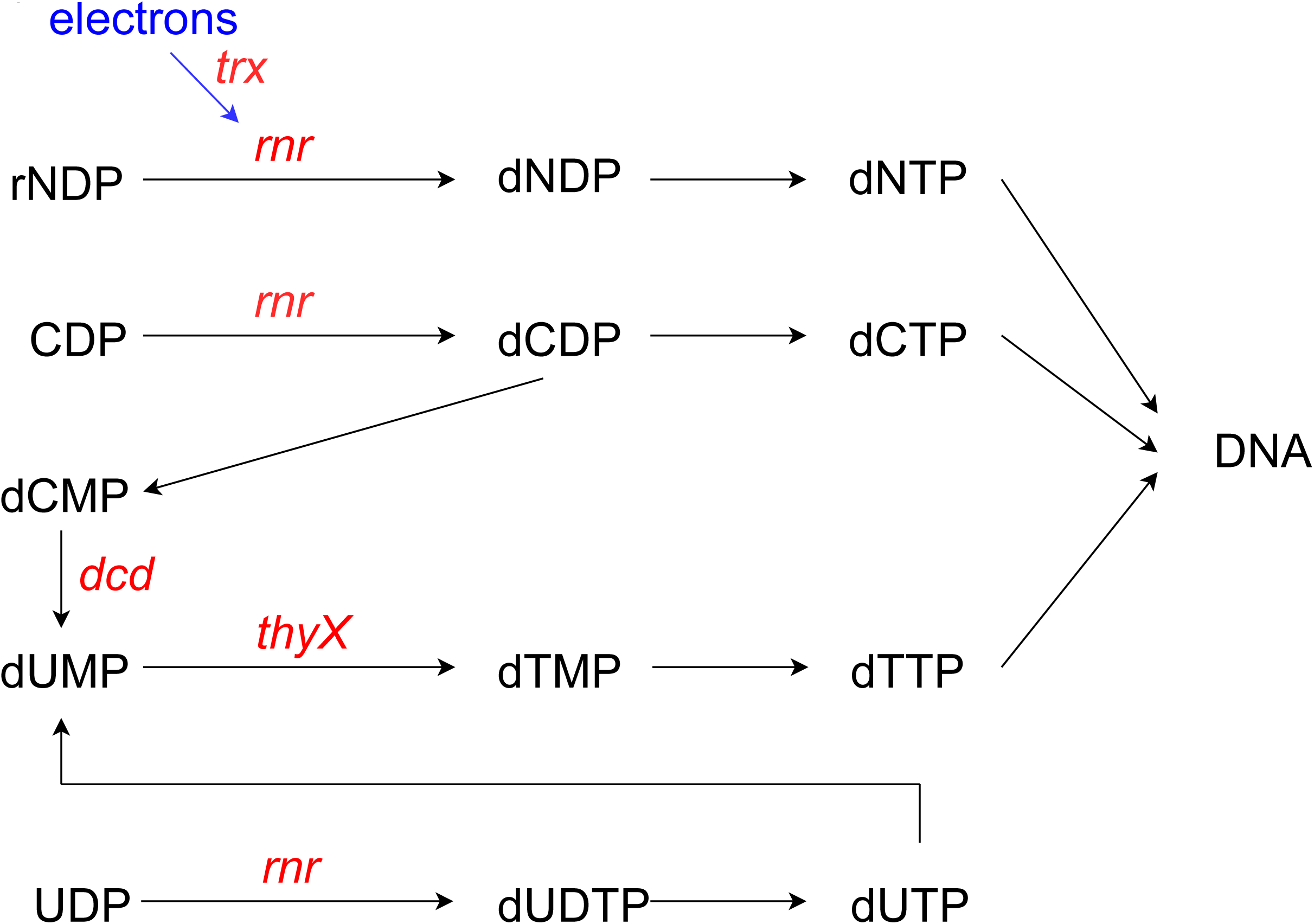
Putative functions of auxiliary metabolic genes in vB_DshP-R7L to promote DNA synthesis in host cells. Phage AMGs are marked in red.

### Origin and evolution of phage vB_DshP-R7L AMGs

To elucidate the evolutionary history of the AMGs in phage vB_DshP-R7L, we identified homologs for each AMG from *Schitoviridae* phages and *Rhodovirinae* roseophages hosts. A gene-sharing network constructed using vConTACT to highlight potential horizontal gene transfer (HGT) events revealed that all auxiliary metabolic genes (AMGs) in vB_DshP-R7L share homologs with *Schitovirinae* phages or *Roseobacteraceae* hosts (M-cytoscapeAMG). Phylogenetic trees for each AMG of vB_DshP-R7L were generated with their homologs from relatives and hosts of *Rhodovirinae* roseophages to further explore their origins (Figure S2). Additionally, the ratio of non-synonymous to synonymous mutation rates (dN/dS) was calculated for each AMG and conserved genes to assess selective pressures.

The *rnr* and *trx* AMGs, conserved within *Rhodovirinae* roseophages, form unique clusters distinct from their homologs. The *rnr* genes of *Rhodovirinae* roseophages grouped with homologs from their *Roseobacteraceae* hosts, suggesting a unique evolutionary path distinct from other *Schitoviridae* subfamilies. Conversely, the *trx* genes of *Rhodovirinae* roseophages clustered with homologs of their hosts and the *Rothmandenesvirinae* subfamily. Indicating a a likely HGT event across different subfamilies of *Schitovirinae* phages. The dN/dS ratios for *rnr* and *trx* in *Rhodovirinae* were 0.12 and 0.14, respectively.

The *thyX* and *dcd* AMGs appear exclusively in four *Rhodovirinae* genera: *Plymouthvirus*, *Baltimorevirus*, *Xianganvirus*, and *Aorunvirus*. Phylogenetic analyses based on concatenate conserved genes suggest these four genera form a monophyletic clade within *Rhodovirinae* subfamily (Figure 2b). The *thyX* genes of *Rhodovirinae* cluster with their homologs of *Rhizobium* azibense and its infecting roseophages (Figure S2). The *dcd* genes of *Rhodovirinae* homologs cluster with homologs from *Cohaesibacter* haloalkalitolerans, *Bradyrhizobium* sp. CCBAU_21365, and phages of the *Erskinevirinae* subfamily. The origins of *thyX* and *dcd* in *Rhodovirinae* likely trace back to an ancient HGT event occurred on the common ancestor of these four genre, with dN/dS ratios of 0.18 and 0.23, respectively.

In *Rhodovirinae*, the distribution of *nanS* genes is limited to RD-1410W1-01 and vB_DshP-R7L, while ribosomal protein genes are found only in vB_DshP-R1 and vB_DshP-R7L. The *nanS* gene sequences are highly similar and cluster with homologs from plasmids of *Roseobacter denitrificans* and its infecting roseophages (Figure S2). Given RD-1410W1-01’s broad host range, including *Roseobacter denitrificans* and *Dinoroseobacter shibae*, the *nanS* gene in vB_DshP-R7L likely originated from a recent HGT event facilitated by RD-1410W1-01 as they both infect *Dinoroseobacter shibae* cell. The ratio of synonymous and nonsynonymous substitutions (dN/dS ratio) for *nanS* among vB_DshP-R7L, RD-1410W1-01, and their host homologs is 0.27. The ribosomal protein genes of *Rhodovirinae* roseophages are genetically similar to their host homologs and significantly different from previously reported viral homologs, indicating they may originate from recent HGT events from their hosts (Figure S3).

We utilized the tRNA adaptation index (tAI) to assess the adaptation of vB_DshP-R7L genes to the phage’s own tRNA pool. The tAI values of vB_DshP-R7L genes ranged from 0.961 to 0.968 when aligned against the viral tRNA pool, and from 0.459 to 0.508 against the *Dinoroseobacter shibae* host tRNA pool. This analysis indicates that the genes of vB_DshP-R7L have evolved to exhibit a high level of adaptation to the tRNAs present within the phage genome, reflecting a significant evolutionary adjustment to optimize translation efficiency in their viral context.

## Discussion

The virion morphology and structural protein profiles of the newly isolated vB_DshP-R7L closely match those reported in previous studies on other *Rhodovirinae* roseophages [23, 25, 27, 68]. Mass spectrometry and electrophoresis analysis identified five conserved structural proteins of *Schitoviridae* family. However, other conserved structural proteins of the *Schitoviridae*, present in the genome of vB_DshP-R7L, were not detected in electrophoretic or mass spectrometry analyses. This discrepancy is likely due to their lower abundance being overshadowed by highly abundant structural proteins, particularly the major capsid protein. Similar challenges in identifying structural proteins have been reported for other *Schitoviridae* phages [27, 69]. Previous studies have indicated that most *Rhodovirinae* roseophages, except for those in the genus *Sanyabayvirus*, can only infect one specific host [23, 27, 70]. Our findings suggest that vB_DshP-R7L exhibits a similar host-specificity, infecting only *Dinoroseobacter shibae* DFL12.

Despite sharing virion structure and host specificity with other *Rhodovirinae* roseophages, vB_DshP-R7L exhibits a unique lytic curve, with a rapid phase lasting five hours—significantly longer than that of its relatives—and a comparatively smaller final burst size [23]. This characteristic likely contributes to the notably small plaque sizes observed, which are too small to measure accurately.

Due to the absence of universally conserved genes in viruses, the International Committee on Taxonomy of Viruses (ICTV) recommends using multiple independent methods for phage classification. These include assessing nucleotide sequence similarity, phylogenetic analysis of conserved genes with appropriate outliers, and the ratio of homologous genes within a viral family [71]. We employed three independent phylogenetic analyses to determine the taxonomic classification of vB_DshP-R7L: genome-wide sequence similarities among all isolated roseophages, concatenated conserved genes among *Schitoviridae* phages, and single marker gene analyses. All results consistently support the classification of vB_DshP-R7L within a novel genus clade of subfamily *Rhodovirinae*. Following the naming tradition for *Rhodovirinae* roseophages, we propose ’*Xianganvirus*’ as the name for this new genus.

The global distribution of *Rhodovirinae* has remained elusive. Zhan *et al*. identified *Rhodovirinae* roseophages in cold bays using pol gene PCR amplification [72], and a previous study reported extremely low relative abundances of *Schitoviridae* phages in marine metagenomic libraries [18]. A recent study highlighted their predominance in estuarine rather than open ocean environments in the Chesapeake Bay and Delaware Bay [73]. Our research attempted to explore the distribution of *Rhodovirinae* roseophages with the newly isolated vB_DshP-R7L, revealed that all detected *Rhodovirinae* roseophages were confined to estuarine ecosystem, specifically within metagenomic samples from the Pearl River Estuary and Delaware Bay. This limited distribution may be due to their restricted host range, potentially hindering their expansion across open oceans.

AMGs are prevalent in marine phages, including those of the *Rhodovirinae* roseophages, where they hijack host metabolism to enhance phage production [39]. In vB_DshP-R7L, AMGs are predominantly located at the 3’ end of the genome, consistent with patterns observed in its close relatives. Six AMGs have been identified in this genome; four are involved in the DNA *de novo* synthesis pathway and may enhance the production of DNA precursors, thus facilitating DNA replication and modification during infection. The remaining two AMGs in vB_DshP-R7L include a gene encoding the ribosomal protein bL12, and another encoding sialate O-acetylesterase (*nanS*). Studies indicate that viral ribosomal protein bL12 genes are typically shorter than their host counterparts, and experiments in vitro have shown that viral encoded bL12 can integrate into the host 70S ribosomes in E. coli [67]. Thus, the bL12 gene in vB_DshP-R7L may enhance viral protein synthesis during infection. Previous research has identified *nanS* genes in the genomes of Stx2a phages infecting E. coli, suggesting that the presence of *nanS* may support phage proliferation in the gastrointestinal microbiome [74, 75]. Given that vB_DshP-R7L was isolated near a populated city and detected in metagenomic samples from the Pearl River Estuary, its *nanS* gene could potentially improve the utilization of carbon and nitrogen sources in estuary environments.

Our analyses indicate that the AMGs of vB_DshP-R7L originate from distinct horizontal gene transfer (HGT) events, each following unique evolutionary trajectories. Phylogenetic analyses of the four DNA synthesis-related AMGs, alongside homologs from other *Rhodovirinae* roseophages, show similar tree topologies to those based on concatenated conserved genes. These genes cluster with homologs from their *Roseobacteraceae* hosts, suggesting ancient HGT events from these hosts to a common ancestor of modern *Rhodovirinae* roseophages, with stable transmission across generations.

Unlike most *Rhodovirinae* roseophages, RD-1410W1-01 has a broad host range. The *nanS* genes from RD-1410W1-01, vB_DshP-R7L form a distinct cluster with homologs from *Roseobacter denitrificans*, which is one reported host of RD-1410W1-01 [22]. This implying RD-1410W1-01 may have facilitated the transfer of the *nanS* gene from *Roseobacter denitrificans* to vB_DshP-R7L, given their overlapping host ranges. The ribosomal protein genes in *Rhodovirinae* roseophages exhibit genetic similarity to their host homologs, and are distinct from other viral homologs. We hypothesis that they have independent evolutionary paths and originate through recent HGT events from their hosts.

Viral AMGs may influence host metabolism during infection, and their ecological roles result in evolutionary pressures, which can be quantified using the ratio of nonsynonymous/synonymous substitution (dN/dS ratio). A gene with dN/dS ratio less than 1 indicates purifying selection, suggesting that the gene is actively eliminating harmful mutations throughout its evolutionary history and could play a crucial role during infection [76]. A gene with dN/dS ratio close to 1 suggests that the gene is predominantly affected by neutral mutations and is likely less critical for viral proliferation. Our results suggest that all AMGs of vB_DshP-R7L exhibit dN/dS ratios significantly below 1, indicating strong purifying selection and implying a vital role in infection processes. Particularly, AMGs involved in the DNA *de novo* synthesis pathway have experienced the highest selection pressures, as evidenced by significantly low dN/dS ratios. The average dN/dS ratio for conserved genes in vB_DshP-R7L is 0.15, and dN/dS ratio of *trx* and rnr genes similar to those of conserved structural genes. AMGs originate from recent HGT events, like *nanS* and ribosomal protein encoding gene, display relatively higher dN/dS values.

Viruses often depend on the host’s tRNA pool and may carry their own tRNA to enhance translation. This exerts evolution pressure on the codon usage of stable viral genes [77]. Our results suggest that all genes in vB_DshP-R7L show a high preference for codon usage aligned with its viral tRNA content, indicating a sophisticated adaptation of the phage genes to the host environment. Despite AMGs being products of HGT, AMGs of vB_DshP-R7L demonstrate a tRNA adaptation index (tAI) similar to that of conserved genes. This similarity emphasizes the substantial selective pressure on AMG codon usage relative to the phage tRNA content, underlining their significant ecological potential.

## Conclusions

In this study, we isolated and characterized a novel roseophage, vB_DshP-R7L, which infects *Roseobacteraceae*, a significant component of coastal bacterial communities. Phylogenetic results reveal that vB_DshP-R7L representing a new genus within the *Rhodovirinae* subfamily of roseophages. Unlike previously characterized *Rhodovirinae* roseophages, vB_DshP-R7L exhibits notably slower growth rates and a reduced burst size. This phage was exclusively detected in metagenomic samples from the Pearl River Estuary, consistent with the environmental specificity observed for all *Rhodovirinae* roseophages to estuarine ecosystems. We identified six AMGs within the genome of vB_DshP-R7L, four are related to *de novo* nucleotide synthesis, while the others may involved in protein production and carbon metabolism. Comparative genomic analyses suggest these AMGs originated from distinct HGT events, and all subject to strong evolution pressure, indicating their stable incorporation into the genome and functional potential in estuarine ecosystems of the South China Sea. Our comprehensive analysis enhances the understanding of the phylogeny and AMG content within the *Rhodovirinae* roseophage clade, offering new insights into the diversity, evolution, and biogeography of *Rhodovirinae* roseophages in marine environments. These findings not only identify a new phage genus but also broaden our knowledge of the phylogenetic diversity and evolutionary dynamics of marine phages.

## Captain of Figures and Tables

Fig. S1 Phylogenetic analysis of isolated *Schitoviridae* phages based on amino acid sequences marker genes. a) Phylogenetic analysis based on vRNAP2 using maximum likelihood method, the bootstrap value is 100. b) Phylogenetic analysis based on major capsid protein using maximum likelihood method, the bootstrap value is 100.

Fig. S2 Phylogenetic trees of each vB_DshP-R7L AMG with homologous from *Schitoviridae* phages and *Roseobacteraceae* hosts. Phage vB_DshP-R7L isolated in this study is highlightened in red, and the background color of each subfamily of *Schitoviridae* on the phylogenetic tree is consistent with their label.

Fig. S3 Comparison of ribosomal protein genes from genomes of *Rhodovirinae* rosepohages and their hosts.

Table S1. Marine metagenomic samples used in this study

## Availability of data and materials

The complete genome and protein sequences have been deposited in GenBank under accession number MZ773648.1 (https://www.ncbi.nlm.nih.gov/nuccore/MZ773648.1). Sequence data for this study are available in the China national center for bioinformation repository under accession number CRA018945 (https://ngdc.cncb.ac.cn/gsa/browse/CRA018945). The genome and protein sequences of vB_DshP-R7L are provided in a GenBank format supplementary material.

## List of abbreviations

AMG: Auxiliary metabolic gene
dN/dS ratio: ratio of synonymous and nonsynonymous substitutions
HGT: horizontal gene transfer
ICTV: International Committee on Taxonomy of Viruses
MRC: marine roseobacter clade
NCBI: National Centre for Biotechnology Information
ORFs: open reading frames
SDS-PAGE: sodium dodecyl sulfate-polyacrylamide gel electrophoresis
tAI: tRNA adaptation index
TEM: Transmission electron microscopy

## Declarations

### Ethics approval and consent to participate

Not applicable.

### Consent for publication

Not applicable.

### Availability of data and materials

The datasets supporting the conclusions of this article are included within the article and its additional files.

### Authors’ contributions

Xingyu Huang wrote the main text of the manuscript and conducted the analysis. Chen Yu was responsible for the purification, while Longfei Lu was responsible for isolation and provided the basic ideas.

## Acknowledgements

Not applicable.

