## Supplementary figures and images for "Isolation and Characterization of a Roseophage Representing a Novel Genus in the N4-like Rhodovirinae Subfamily Distributed in Estuarine Waters"

### FigureS1.pdf

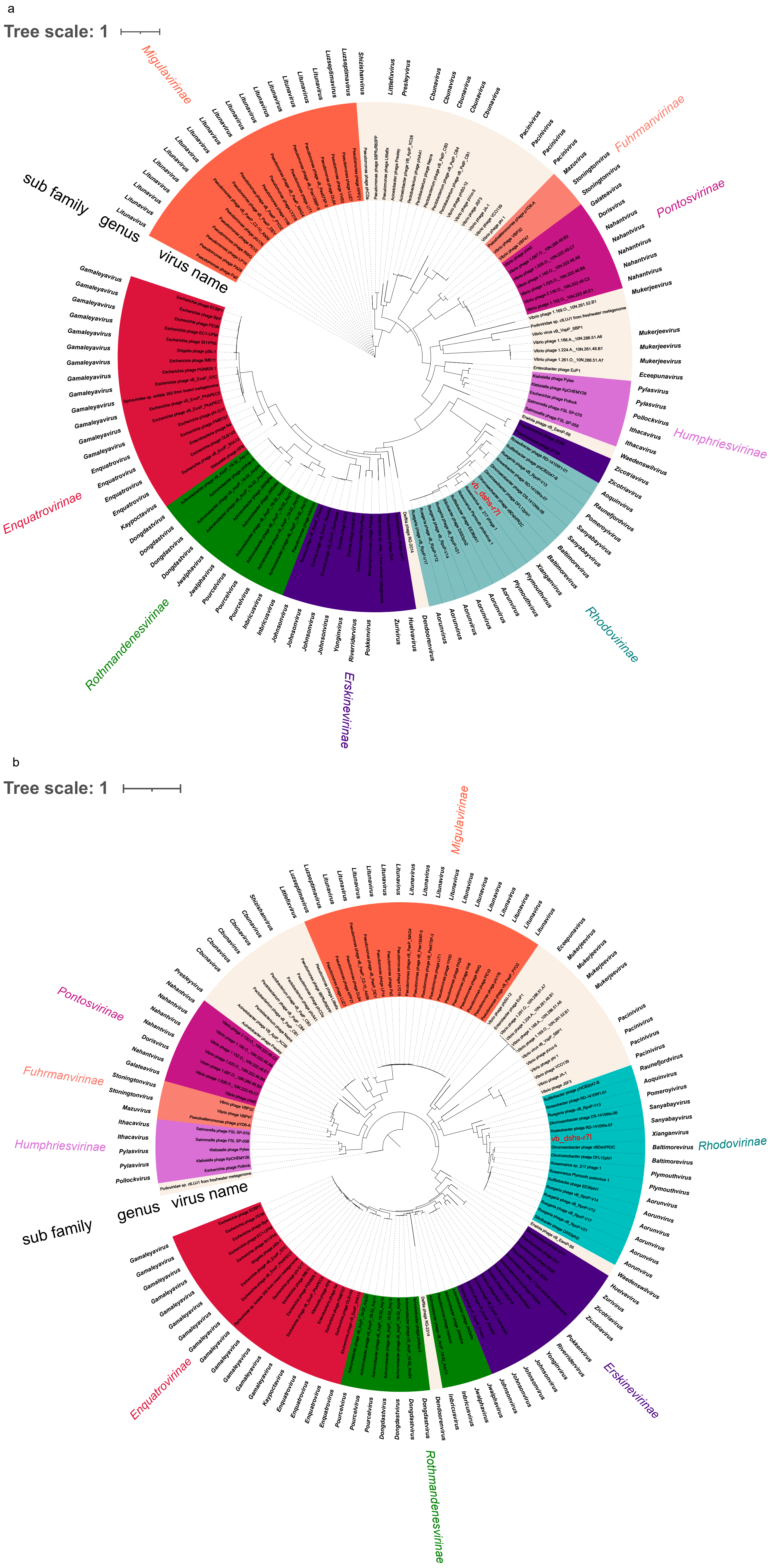

### FigureS2.pdf

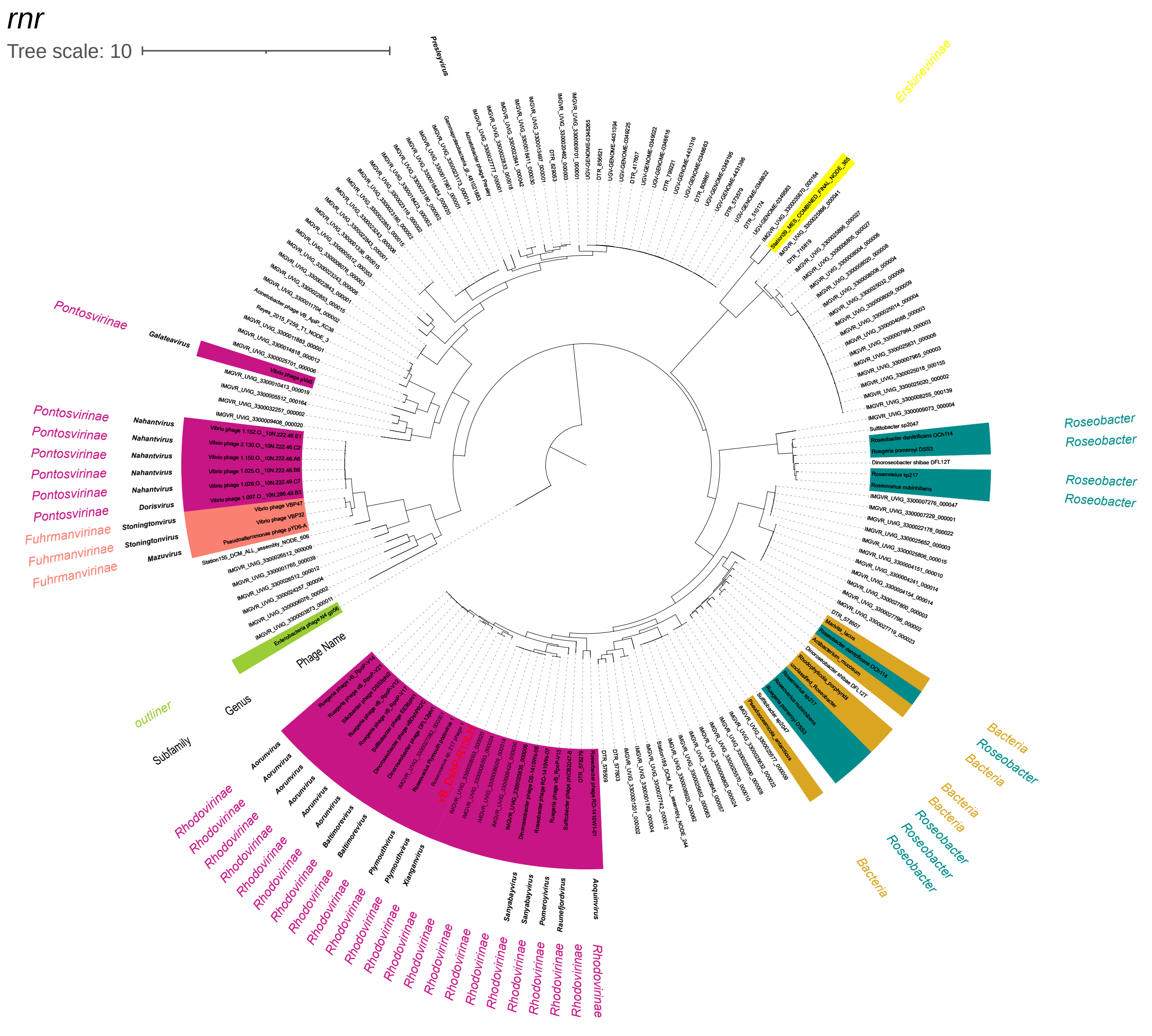

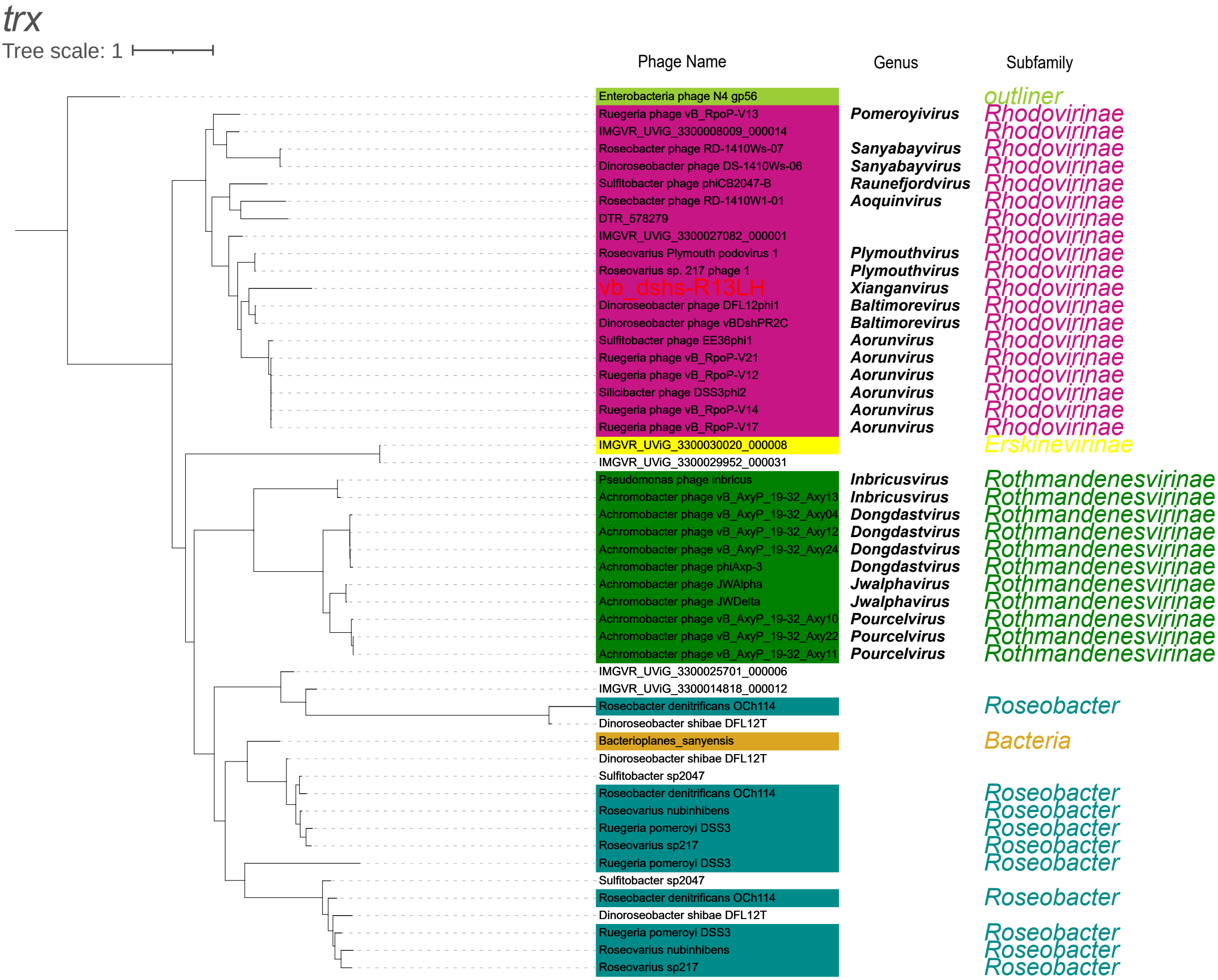

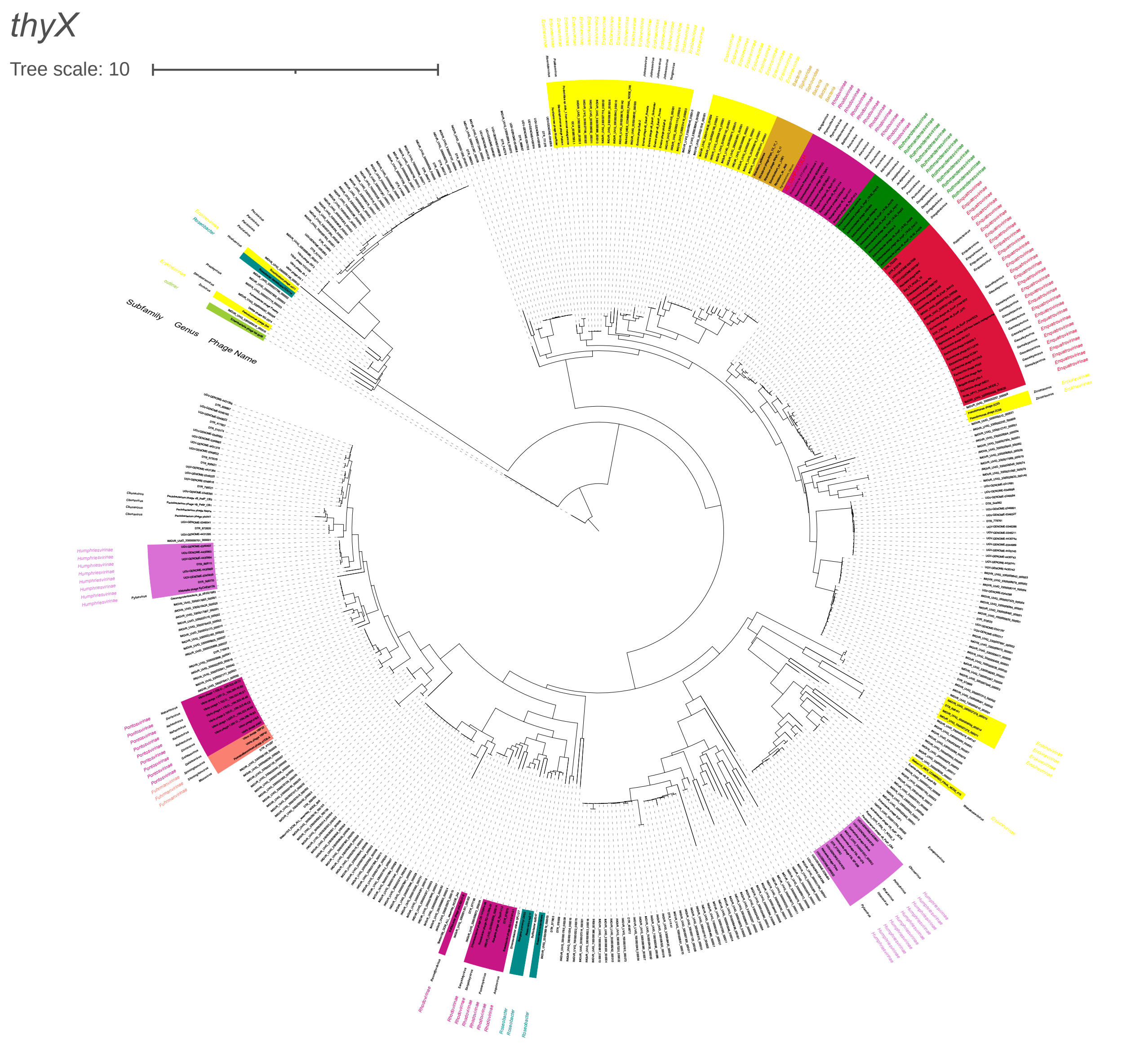

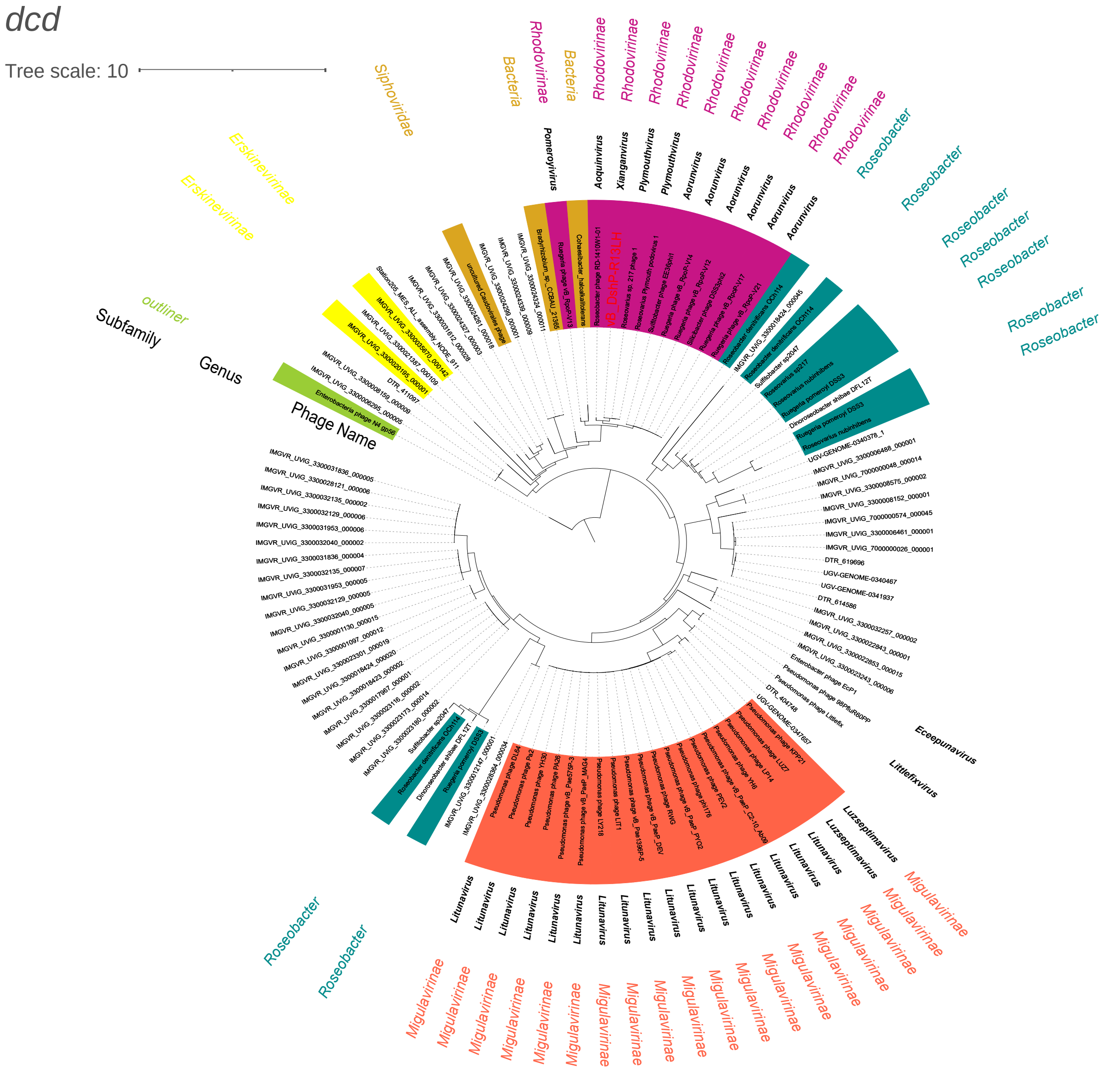

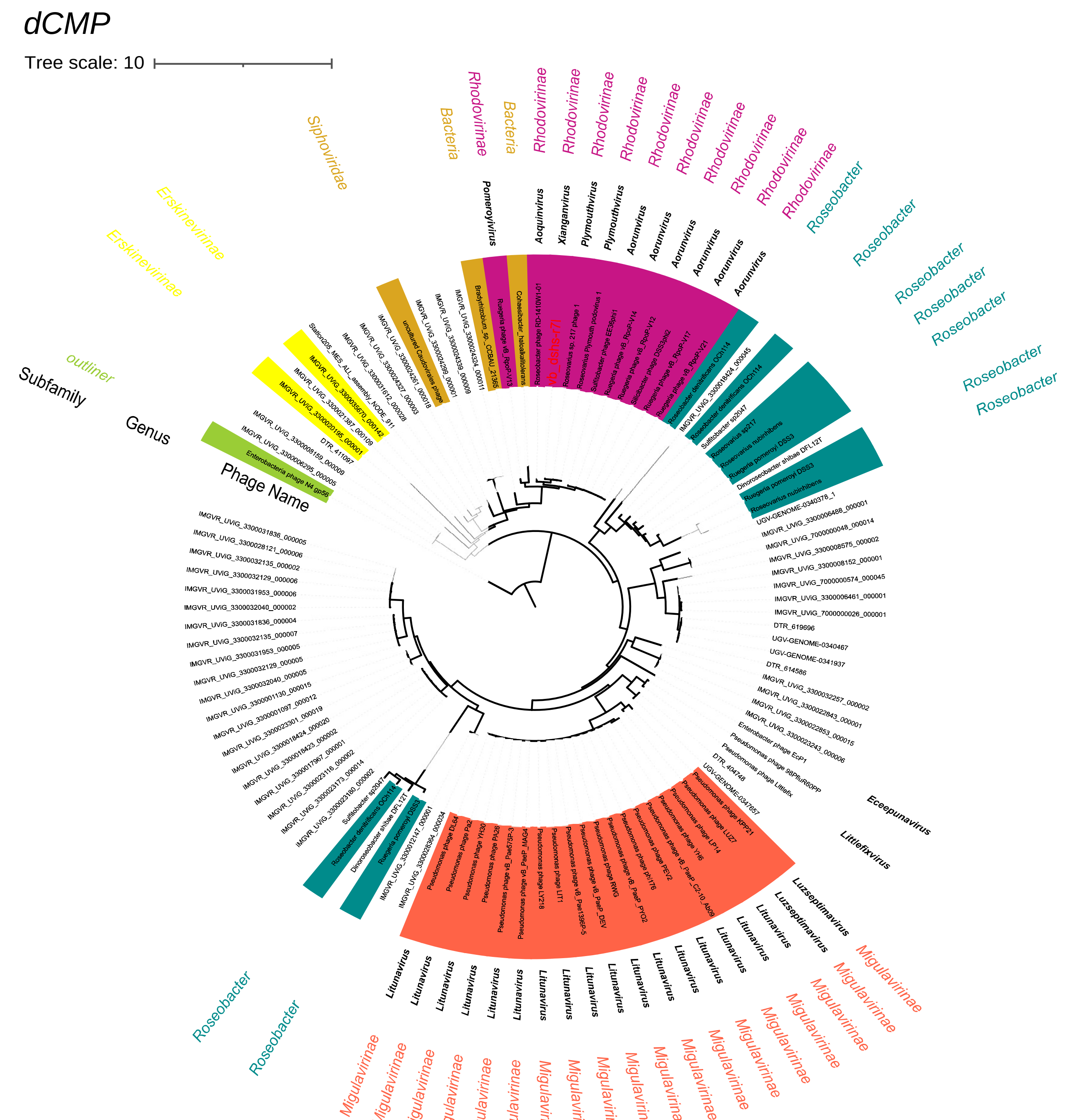

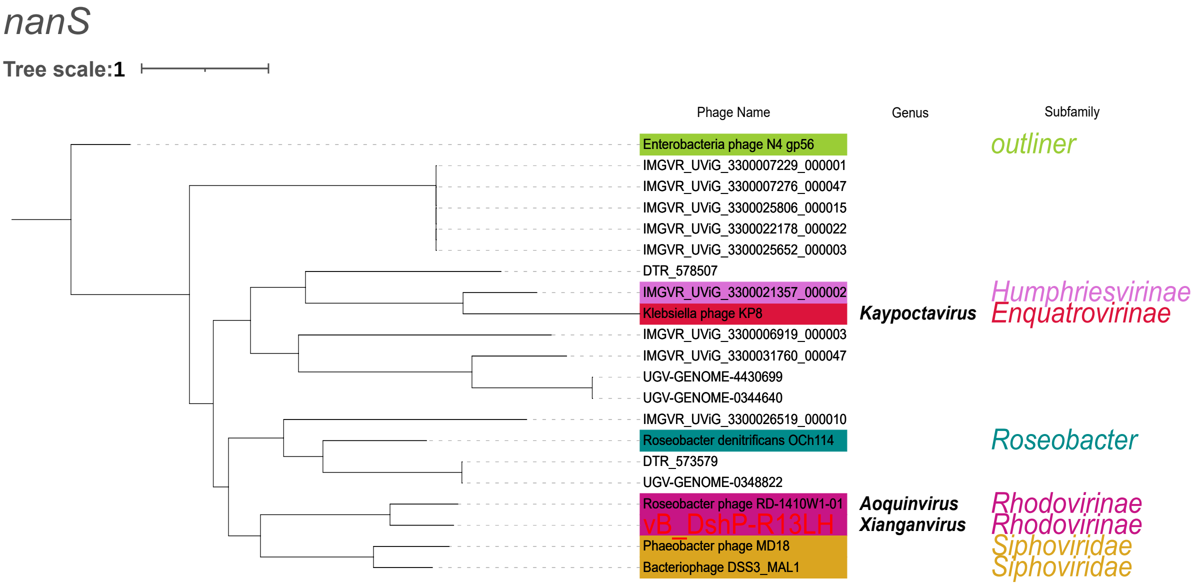

### FigureS3.pdf

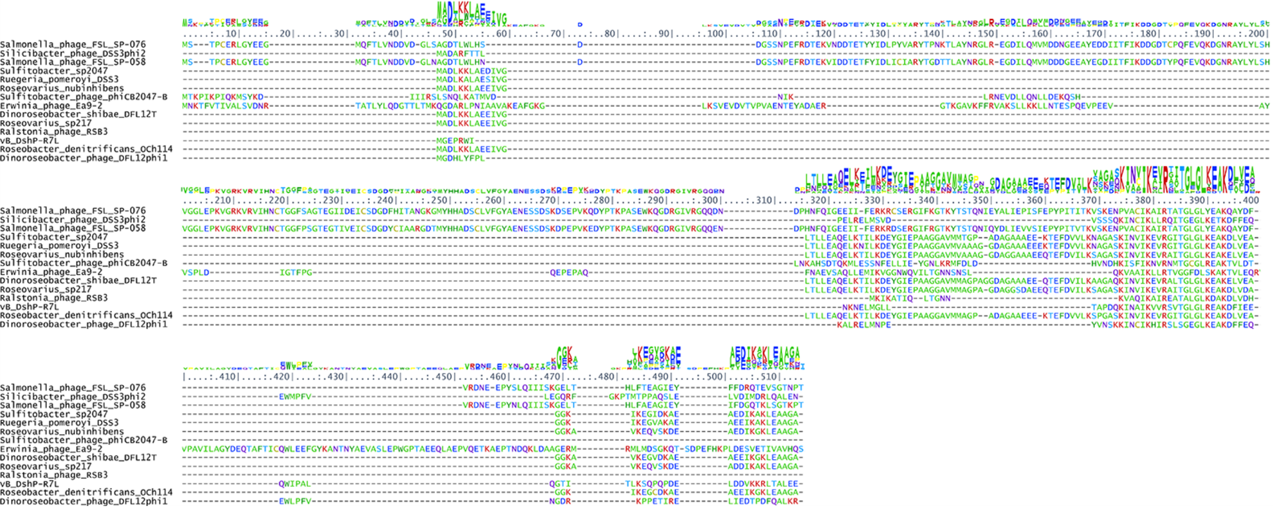
